# A metal ion-dependent mechanism of RAD51 nucleoprotein filament disassembly

**DOI:** 10.1101/2023.02.18.529066

**Authors:** Robert Appleby, Daniel Bollschweiler, Dimitri Y. Chirgadze, Luay Joudeh, Luca Pellegrini

**Author notes:** Max Planck Institute of Biochemistry, CryoEM facility, Am Klopferspitz 18, 82152 Planegg, Germany (Daniel Bollschweiler).

## Abstract

The RAD51 ATPase polymerises on single-stranded DNA to form nucleoprotein filaments (NPFs) that are critical intermediates in the DNA strand-exchange reactions of Homologous Recombination (HR). ATP binding is important to maintain the NPF in a competent conformation for strand pairing and exchange. Once strand exchange is completed, ATP hydrolysis licenses the filament for disassembly. Here we show using high-resolution cryoEM that the ATP-binding site of the RAD51 NPF contains a second metal ion. In the presence of ATP, the metal ion promotes the local folding of RAD51 into the conformation required for DNA binding. The metal ion is absent in the structure of an ADP-bound RAD51 filament, that rearranges in a conformation incompatible with DNA binding. The presence of the second metal ion explains how RAD51 couples the nucleotide state of the filament to DNA binding. We propose that loss of the second metal ion upon ATP hydrolysis drives RAD51 dissociation from the DNA and weakens filament stability, thus contributing to NPF disassembly, a critical step in the completion of HR.

## Introduction

Maintenance of genomic stability in bacteria, archaea and eukaryotes depends critically on the ability to exchange strands between homologous DNA molecules in a reaction known as Homologous Recombination (HR). The steps of HR comprise the invasion of a target duplex DNA by a single-stranded (ss) DNA segment, followed by search for sequence homology, strand pairing between complementary DNA sequences and strand exchange (Filippo et al., 2008; Prakash et al., 2015). Different flavours of HR intervene during various processes of nucleic acid metabolism, such as double-strand break repair (Jasin & Rothstein, 2013; Wright et al., 2018), DNA replication (Saada et al., 2018; Tye et al., 2020) and the crossover reaction of meiotic prophase (Sansam & Pezza, 2015; Zickler & Kleckner, 2015).

A highly conserved super-family of ATPases – including eukaryotic RAD51, archaeal RadA and bacterial RecA as representative members – catalyses the strand-exchange reactions of HR. These strand-exchange proteins (recombinases) polymerise on the single-stranded DNA generated from the controlled resection of DNA ends, forming a helical nucleoprotein filament (NPF). The NPF executes the strand invasion of the target double-stranded (ds) DNA, to form a three-strand intermediate (synapsis), within which the search for homology and strand exchange take place. Given its central role in DNA metabolism and genomic stability, the biochemical reaction of HR has been the focus of intense research efforts over the past 30 years.

Earlier X-ray crystallography and electron microscopy studies of HR recombinases elucidated the helical features of their filamentous structures, demonstrating that bound DNA was both stretched and underwound (Capua et al., 1982; Conway et al., 2004; Ogawa et al., 1993; Story et al., 1992; Yu et al., 2001). The crystallographic studies culminated in the determination of the NPF structure of bacterial RecA, which illustrated how the phosphodiester backbone of the ssDNA is embedded within the protein sheath of the polymeric recombinase, whilst the bases project outwards towards the solvent, poised for the homology search (Chen et al., 2008). The recent development of modern cryoEM approaches has made it easier to image at high resolution the interaction of eukaryotic RAD51 with the DNA within the NPF (Xu et al., 2017). These studies showed that the RecA and RAD51 filaments adopt a common mechanism of DNA interaction; the L1 and L2 loops protruding from adjacent ATPase domains grip the DNA strand, unstacking the bases at each protomer interface and partitioning the nucleotides into triplets.

ATP binding and hydrolysis are essential for RAD51 function. ATP binds at the protomer-protomer interface in the RAD51 filament and promotes the adoption of a filament conformation that is competent for the strand pairing and exchange reactions (Xu et al., 2017). ATP hydrolysis is not required for activity *in vitro* as RAD51’s Walker A mutants defective in ATP hydrolysis can mediate strand exchange (Chi et al., 2006; Sung & Stratton, 1996); moreover, inhibition of ATP hydrolysis using Ca^2+^ or a non-hydrolysable ATP analogue potentiates strand-exchange activity by keeping the filament in a competent state for recombination (Bugreev & Mazin, 2004). However, the ability to hydrolyse ATP is clearly critical *in vivo* (Morgan et al., 2002; Morrison et al., 1999; Stark et al., 2004), pointing to its importance in the completion of Homologous Recombination in cells. Biochemical evidence shows that that ATP hydrolysis causes RAD51 release from DNA and is necessary for filament disassembly (Li et al., 2007; Mameren et al., 2009; Namsaraev & Berg, 1998; Ristic et al., 2005): the requirement for ATP hydrolysis is therefore explained by the need to remove RAD51 from DNA once recombination is completed. Dissociation of RAD51 protomers is thought to proceed from the ends of the filament by RAD51 molecules that have hydrolysed their ATP to ADP (Mameren et al., 2009). As ATP hydrolysis happens randomly throughout the filament, disassembly takes place by bursts of ADP-bound RAD51 molecules at the filament end, until the next ATP-bound RAD51 becomes terminal (Mameren et al., 2009).

The consequence of ATP hydrolysis on the RAD51 NPF structure and why it should licence its disassembly is unclear. Crystallographic analysis of the archaeal RAD51 orthologue RadA bound to the non-hydrolysable ATP-homologue AMPPNP had shown the presence of a potassium ion at the ATP-binding site, in contact with the gamma phosphate and making bridging interactions with the neighbouring RadA protomer in the crystallographic filament (Wu et al., 2004, 2005). The position and range of interactions of the potassium ion suggested a role for it in organising the L2 loop of the neighbouring RadA molecule in a conformation competent for DNA binding, in agreement with the known stimulatory effects of monovalent cations on DNA binding and strand-exchange activity (K.-S. Shim et al., 2006; S Sigurdsson et al., 2001).

Here we used high-resolution cryoEM to investigate the possible presence and functional role of a second metal ion in pre- and post-synaptic RAD51 NPFs. We find that the ATP-binding site contains a second metal ion, in addition to the canonical ion bound within the Walker A motif, in analogous position to the potassium ion observed in the crystal structure of RadA. We further show that loss of the second metal ion in the cryoEM structure of the ADP-bound RAD51 filament leads to a conformational rearrangement in the L2 loop that renders it incompatible with DNA binding. Thus, ATP hydrolysis drives a conformational transition of the filament to a state that is unable to bind DNA and weakens RAD51 self-association, licensing the NPF for disassembly.

## RESULTS

### A second metal ion at the ATP site of the pre-synaptic RAD51 filament

We determined the cryoEM structure of human RAD51 bound to a ssDNA 60mer and Ca^2+^ATP at 3.8 Å resolution (Figure 1A, Supplementary figure 1, Supplementary table 1, Methods). Inspection of the density map at the ATP-binding site showed the presence of a peak near the gamma phosphate and at the interface with the neighbouring RAD51 protomer, which was not accounted for by the known features of the ATP ligand and its protein-binding moieties (Figure 1B). In addition to the gamma phosphate, the density peak was surrounded by the main-chain carbonyl groups of A293, H294 and S296 and by the side-chain carboxylate of D316 in the ATPase domain of the adjacent RAD51 protomer (Figure 1C). The position of the peak was analogous to that of the potassium ion bound in the crystal structure of archaeal RadA (Wu et al., 2005) (Figure 1D). The presence of negatively charged oxygen moieties in suitable positions to act as ligands for a divalent metal ion supported the presence of a previously unidentified Ca^2+^ ion bound to ATP, in addition to the canonical Ca^2+^ ion bound within the Walker A motif and required for ATP hydrolysis.

**Figure 1.**
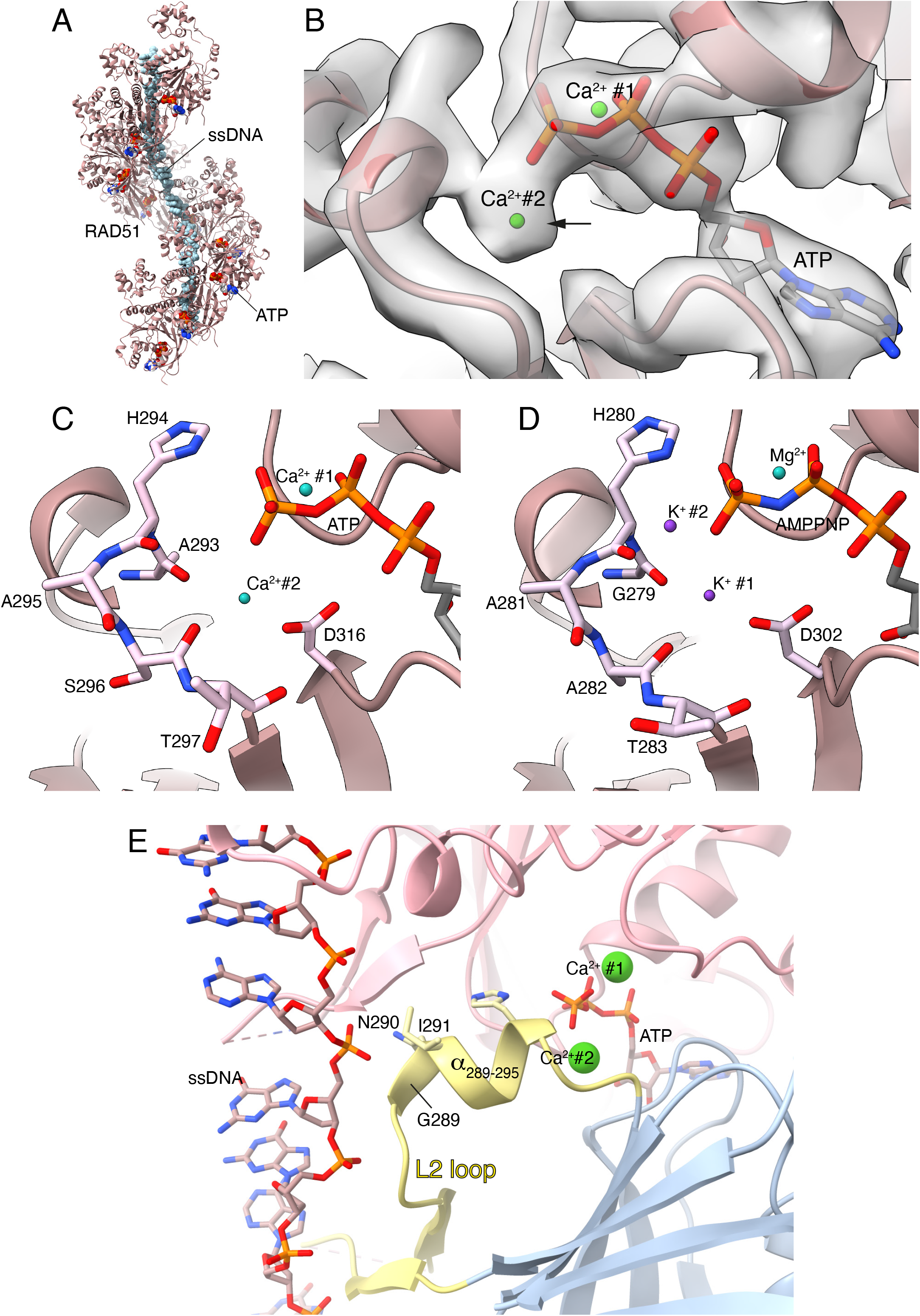
The second metal cation in the pre-synaptic RAD51 filament. **A** CryoEM structure of the human RAD51-ssDNA-ATP filament. RAD51 is shown as ribbons, ssDNA and ATP as spacefill models. **B** CryoEM map at the ATP site, with the density peak for the second Ca^2+^ ion marked by an arrow. The two Ca^2+^ cations are drawn as green spheres. **C** Molecular details of the interactions of the second Ca^2+^ cation with ATP and with main-chain and side-chain atoms of surrounding RAD51 residues. The two Ca^2+^ cations are drawn as green spheres. The amino acids that interact with Ca^2+^ and ATP are shown in stick representation. The rest of the RAD51 is drawn as a light brown ribbon. **D** Molecular details of the interactions of the two K^+^ cations with AMPPNP and surrounding main-chain and side chain residues of the archaeal RAD51-orthologue RadA (PDB ID 2FPM). The protein and ATP are drawn as in panel C, the two K^+^ cations are shown as purple spheres, the Mg^2+^ cation in green. **E** The RAD51 *α*-helix spanning residues 289 to 295 of the L2 loop couples DNA binding to ATP. The second Ca^2+^ cation stabilises the helical conformation of L2 *α*_289-295_ by interacting with the C-terminal end of the helix. The two RAD51 protomers are shown as ribbons in pink and light blue. The conformation of the L2 loop is highlighted in pale yellow, with the two Ca^2+^ cations drawn as green spheres. The ssDNA is drawn in stick representation and coloured according to atom type. The side chains of amino acids G289, N290, I291 at the N-end of the L2 *α*_289-295_ and of H294 at the C-end are drawn explicitly.

The structure showed that the second metal ion contacts simultaneously ATP and the DNA-binding L2 loop in the adjacent RAD51 protomer (Figure 1E). Thus, its location allows it to fulfil concurrently three related functions: to sense the hydrolysis state of the filament by contacting the gamma phosphate, to modulate the affinity of RAD51 for DNA by affecting the conformation of the L2 loop and to strengthen the association between proximal RAD51 molecules in the filament. Together, these interactions provide the basis for a metal-dependent mechanism of coupling the hydrolysis state of the nucleotide to the filament’s ability to bind DNA. By interacting with the carbonyl moieties of residues A293, H294 and S296, the metal ion helps nucleate the folding of the short alpha helix spanning residues G289 to A295 (*α*_289-295_) that anchors the L2 loop to the core of the RAD51 ATPase domain (Figure 1E). This alpha helix has a crucial role in linking DNA binding to the ATP moiety: at its N-end, G289, N290 and I291 pack against the phosphate backbone of the DNA while at its C-end the imidazole ring in the side chain of H294 hydrogen bonds to the gamma phosphate.

These results show that the second metal ion promotes the active state of the RAD51 filament, by inducing the ATP-dependent conformation of the L2 loop that is required for DNA binding.

### The second metal ion is present in the post-synaptic RAD51 filament

To determine whether the second metal ion is present in a RAD51 filament containing dsDNA, which is thought to represent the post-synaptic state of the filament after completion of strand-exchange, we examined the 2.9 Å cryoEM structure of human RAD51 bound to dsDNA and ATP in the same Ca^2+^ buffer (Figure 2A, Supplementary figure 2, Supplementary table 1, Methods). To capture a bona fide post-synaptic filament state, we incubated RAD51 with ssDNA and a dsDNA that contained a central mismatched region with perfect complementarity for the ssDNA (see Methods). The resulting high-resolution structure showed clear, continuous density for dsDNA, as expected for a post-synaptic filament.

**Figure 2.**
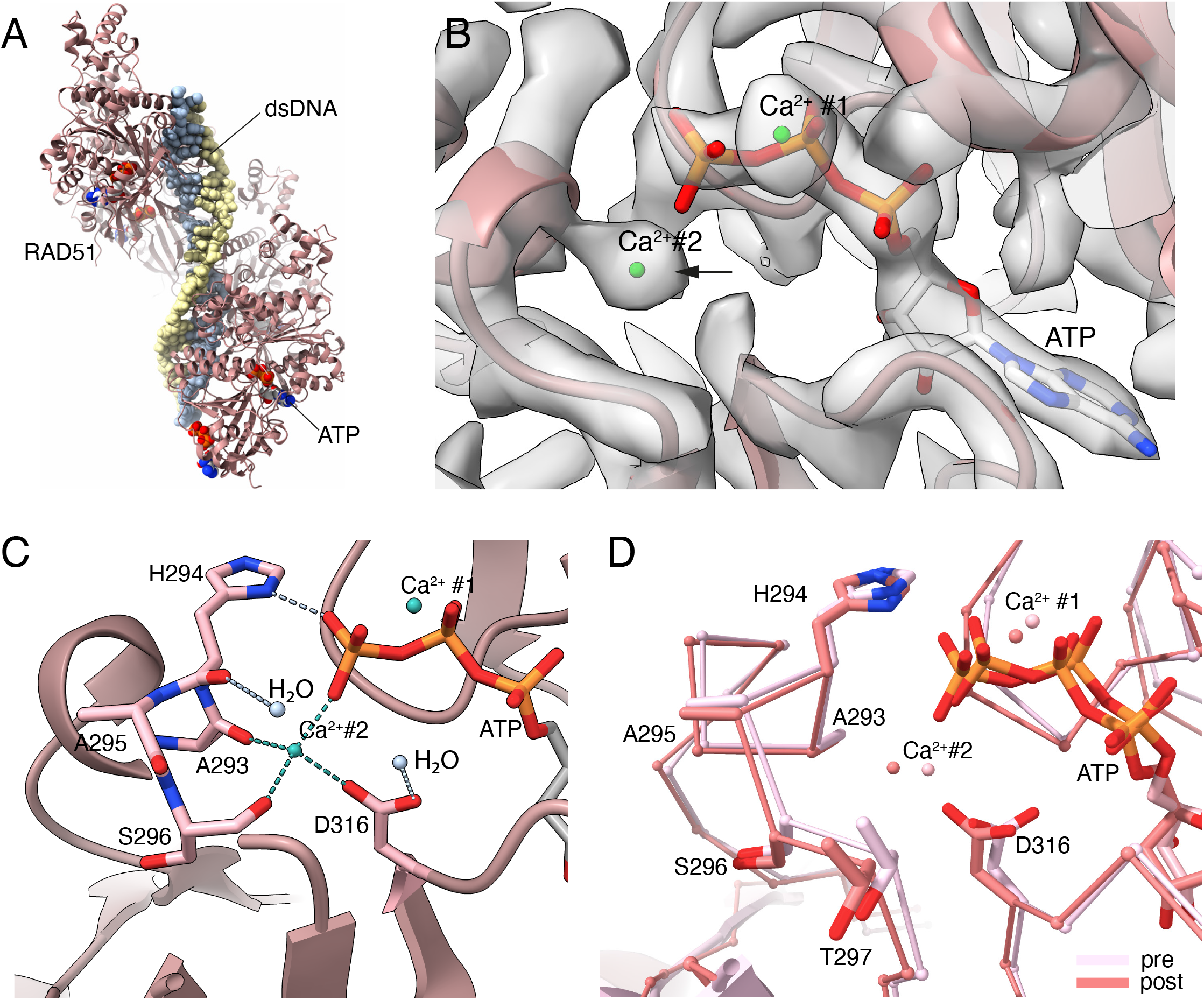
The second metal cation in the post-synaptic RAD51 filament. **A** CryoEM structure of the human RAD51-dsDNA-ATP filament. RAD51 is shown as ribbons, dsDNA and ATP as spacefill models. **B** CryoEM map at the ATP site, with the density peak for the second Ca^2+^ ion marked by an arrow. The two Ca^2+^ cations are drawn as green spheres. **C** Molecular details of the interactions of the second Ca^2+^ cation with ATP and the main-chain and side-chain atoms of surrounding RAD51 residues. The two Ca^2+^ cations are drawn as green spheres. The amino acids that interact with Ca^2+^ and ATP are shown in stick representation. The rest of the RAD51 is drawn as a light brown ribbon. The interactions of the second Ca^2+^ cation with the gamma phosphate of ATP, the carbonyl groups of A293, S296 and the carboxylate group of D316 are highlighted as green dashed lines. The hydrogen bond of the H294 sidechain imidazole with the gamma phosphate and of two water molecules with the main-chain carbonyl group of H294 and the carboxylate of D316 are also shown. **D** Superposition of pre- and post-synaptic filament structures, illustrating the high degree of similarity at the ATP binding site. RAD51 is shown as an alpha carbon trace and relevant side chains are drawn explicitly in stick representation. The pre-synaptic structure is drawn in light pink and the post-synaptic structure in darker pink.

The structure of the post-synaptic NPF was highly similar to that of the pre-synaptic NPF, in agreement with what previously reported (Xu et al., 2017). Inspection of the ATP binding site revealed an unexplained density peak at the same position as in the pre-synaptic filament (Figure 2B), indicating the presence of a bound metal ion that was engaged in the same set of interactions as observed in the pre-synaptic filament structure. The higher resolution of the post-synaptic structure confirmed the gamma phosphate of ATP, the carbonyl groups of A293, S296 and the side-chain carboxylate of D316 of the adjacent RAD51 molecule as the coordinating oxygen atoms (Figure 2C). This apparent tetrahedral coordination is likely to be completed with water molecules to give a more common bi-pyramidal coordination, which would also be the favoured coordination of a physiological Mg^2+^ ion.

These findings show that the second metal ion is present and bound in the same fashion to both pre- and post-synaptic filaments (Figure 2D). It is therefore likely that the second metal ion is present throughout the reactions of homology search and strand exchange, to maintain the L2 loop in the required conformation for DNA binding.

### Structure of the ADP-bound filament

As the metal ion is directly coordinated by the gamma phosphate of ATP, a prediction of our findings is that it will be lost once ATP is hydrolysed to ADP. We prepared a RAD51 NPF sample in the presence of dsDNA and ADP in Ca^2+^ buffer, to mimic a post-synaptic filament that had hydrolysed ATP and determined its cryoEM structure at 3.6 Å resolution (Figure 3A, Supplementary figure 3, Supplementary table 1, Methods). 3D classification showed that, relative to the pre- and post-synaptic NPFs, the ADP-bound filament displayed an increase in protomer rise and a decrease in twist angle. Inspection of the map at the ATP binding site showed the presence of the canonical metal ion in the Walker A motif but no density peak for the second metal ion (Figure 3B), as expected because of the missing gamma phosphate.

**Figure 3.**
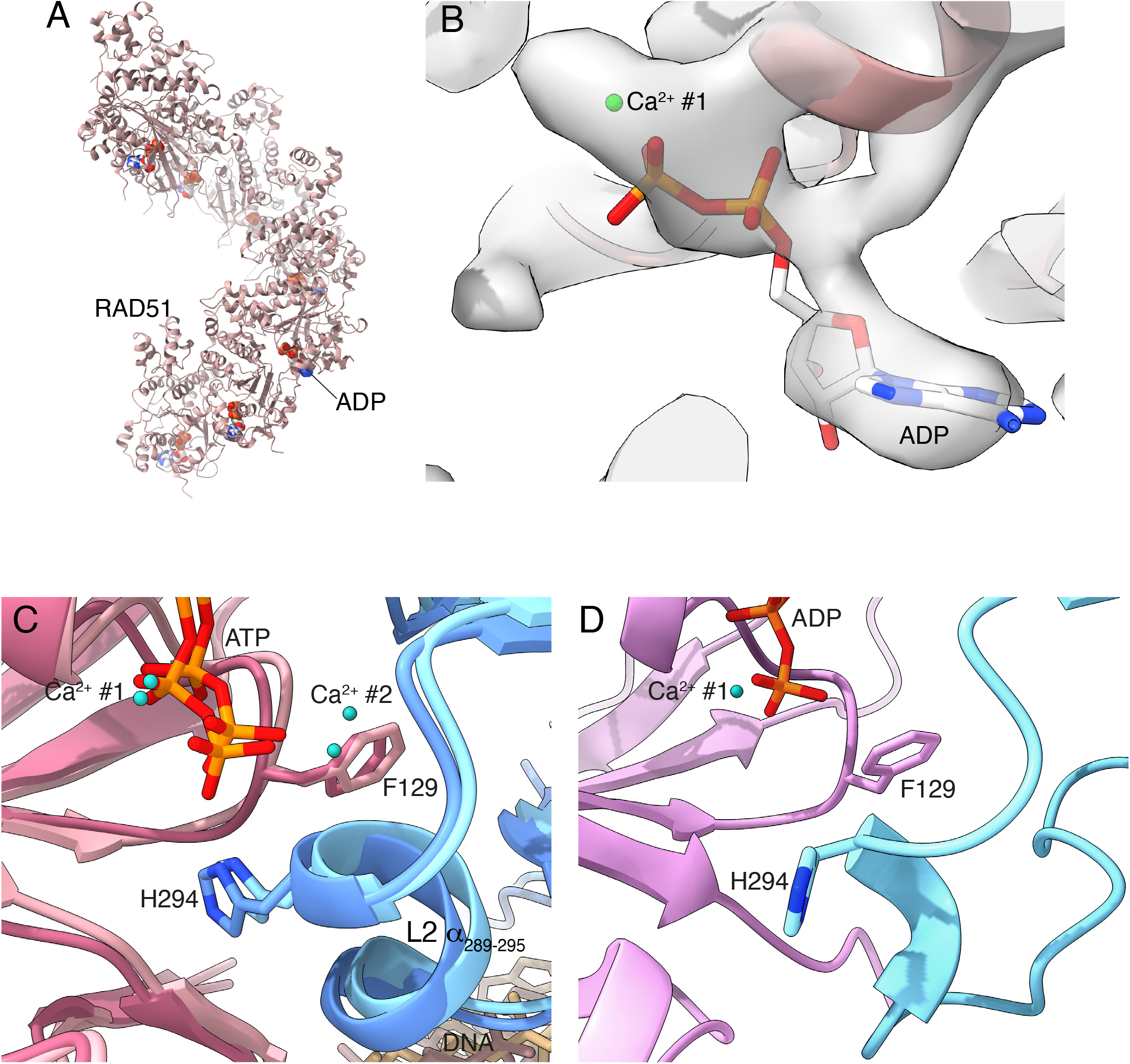
CryoEM structure of the RAD51-ADP filament. **A** Ribbon representation of the ADP-bound RAD51 filament. **B** CryoEM map at the ADP site. The position of the Ca^2+^ cation bound within the Walker A motif is shown as a green sphere. Panels **C** and **D** show the side-chain conformations of F129 in the L2 loop in the presence of ATP and ADP, respectively. In the pre- and post-synaptic filaments (panel C), the F129 sidechain packs against the alpha helix spanning residues 289-295, helping to stabilise the L2 conformation that is conducive to DNA binding. In the ADP-bound filament (panel D), the F129 side chain adopts a rotameric position made accessible by the lack of the gamma phosphate and second metal ion, and that contributes to altering the L2 conformation. Panels C and D are drawn in the same way, with the RAD51 protomers shown as pink and blue ribbons, the side chains of F129, H294, ATP and ADP drawn explicitly in stick representation, and the Ca^2+^ cations as green spheres.

The structure further showed that the conformation of the L2 loop had drastically altered. The side chain of F129 in the Walker A motif, which packs against A293 of the adjacent protomer in the ATP-bound filament, had rotated to partially fill the gap left by the missing gamma phosphate (Figure 3C, D). The steric hindrance of the new F129 rotamer caused the helical residues at the C-end of the L2 loop to swing away and partially unfold. Surprisingly, rather than becoming disordered, the L2 loop transitioned to a compact random-coil conformation (Figure 4), anchored by hydrophobic interactions of I287, I291, I292 and P286 to the ATPase domain (Supplementary figure 4A). The new L2 trajectory appears to be incompatible with DNA binding due to steric clashes with the phosphoester backbone (Supplementary figure 4B), and in fact no density for DNA was observed in any of the 3D classes. We surmise that DNA must have dissociated from RAD51 during incubation prior to grid freezing as a result of the conformational change in L2. It is also possible that the DNA remains weakly associated with the filament in a range of conformations that are not detectable by cryoEM.

**Figure 4.**
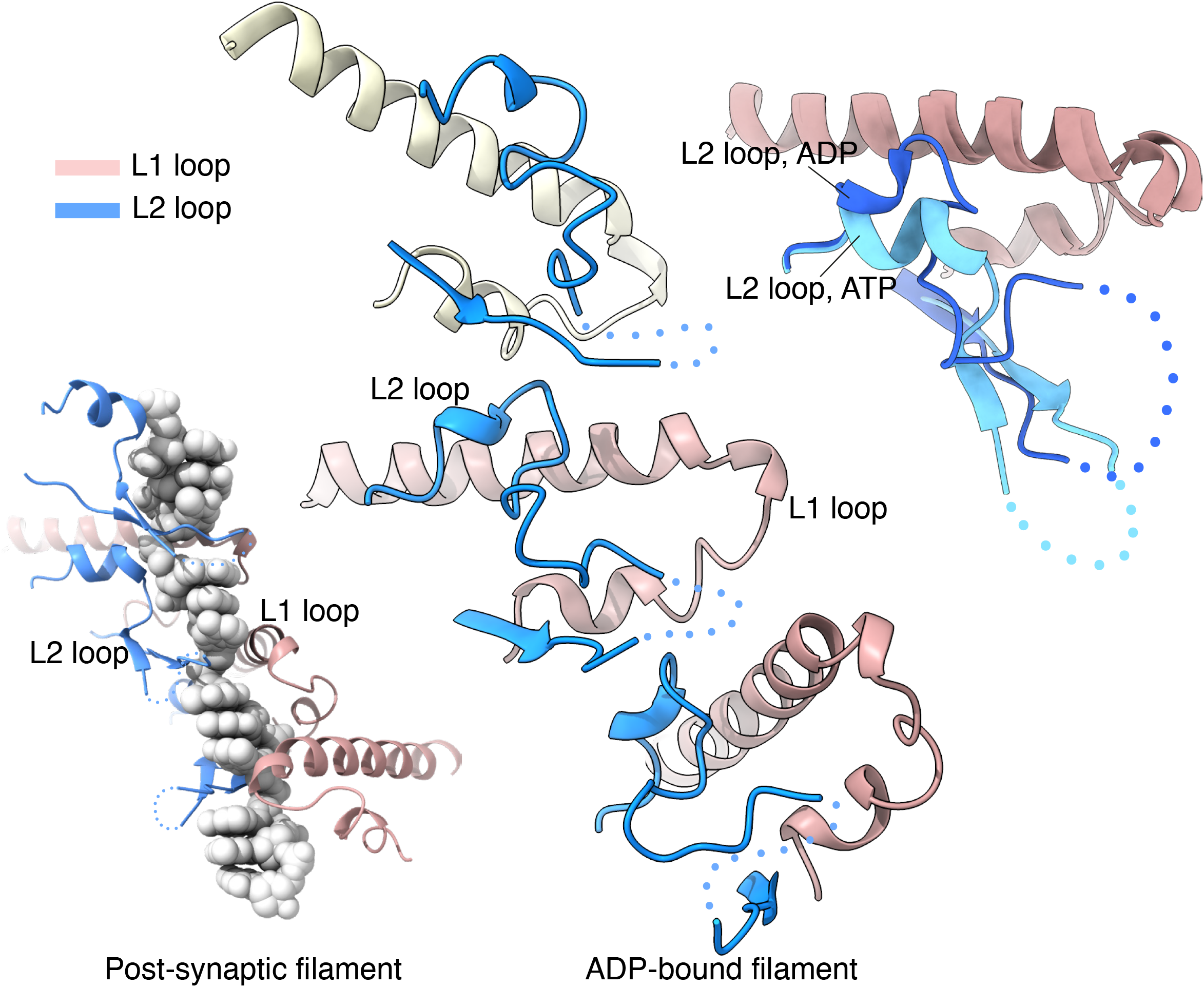
Comparison of the L1 and L2 loop conformations in the RAD51-ADP and post-synaptic filaments. In the central panel, the L1 and L2 loops for three adjacent protomers of the ADP-bound RAD51 filament are shown, drawn as light brown (L1) and light blue (L2) ribbons. For comparison, the arrangement of the L1 and L2 loops in the structure of the post-synaptic filament is shown in the bottom left panel, coloured in the same way; only one strand of DNA is shown, as spacefill model. The top right panel shows a superposition of the L1 and L2 loops in the post-synaptic and ADP filaments, to highlight the different conformation of the L2 loop in the two structures. The L2 loop is in darker blue in the ADP-bound filament and in lighter blue in the post-synaptic filament, whereas the L1 loop is shown in light brown. In the three panels, the missing segment at the tip of the L2 loop is drawn as dots.

These finding show that the presence of the second metal ion depends on the hydrolysis state of the bound nucleotide, and that ATP hydrolysis promotes a conformational change in the L2 loop, with ensuing loss of DNA binding. Thus, the structure of the ADP-bound filament identifies an intermediate state of the filament, where DNA has dissociated and the filament appears poised for disassembly.

## DISCUSSION

The presence of a second metal ion in RAD51 filaments had not been reported before, even though cryoEM structures of pre- and post-synaptic RAD51 NPFs have been published(Xu et al., 2017). It is likely that the resolution of these structures was insufficient for the identification of the metal ion in the map. However, we found clear evidence for it in a recent cryoEM map of a pre-synaptic RAD51 filament at 2.97 Å (EMD 31158, PDB 7EJC) (Xu et al., 2021), and in reconstructions of pre- and post-synaptic filaments of RAD51’s meiotic orthologue DMC1 (pre-synaptic: EMD 30311, PDB 7C9C at 3.33 Å; post-synaptic: EMD 31154, PDB 7EJ7 at 3.41 Å) (Luo et al., 2021) in agreement with our observation, although the authors did not discuss it (Supplementary figure 5).

**Figure 5.**
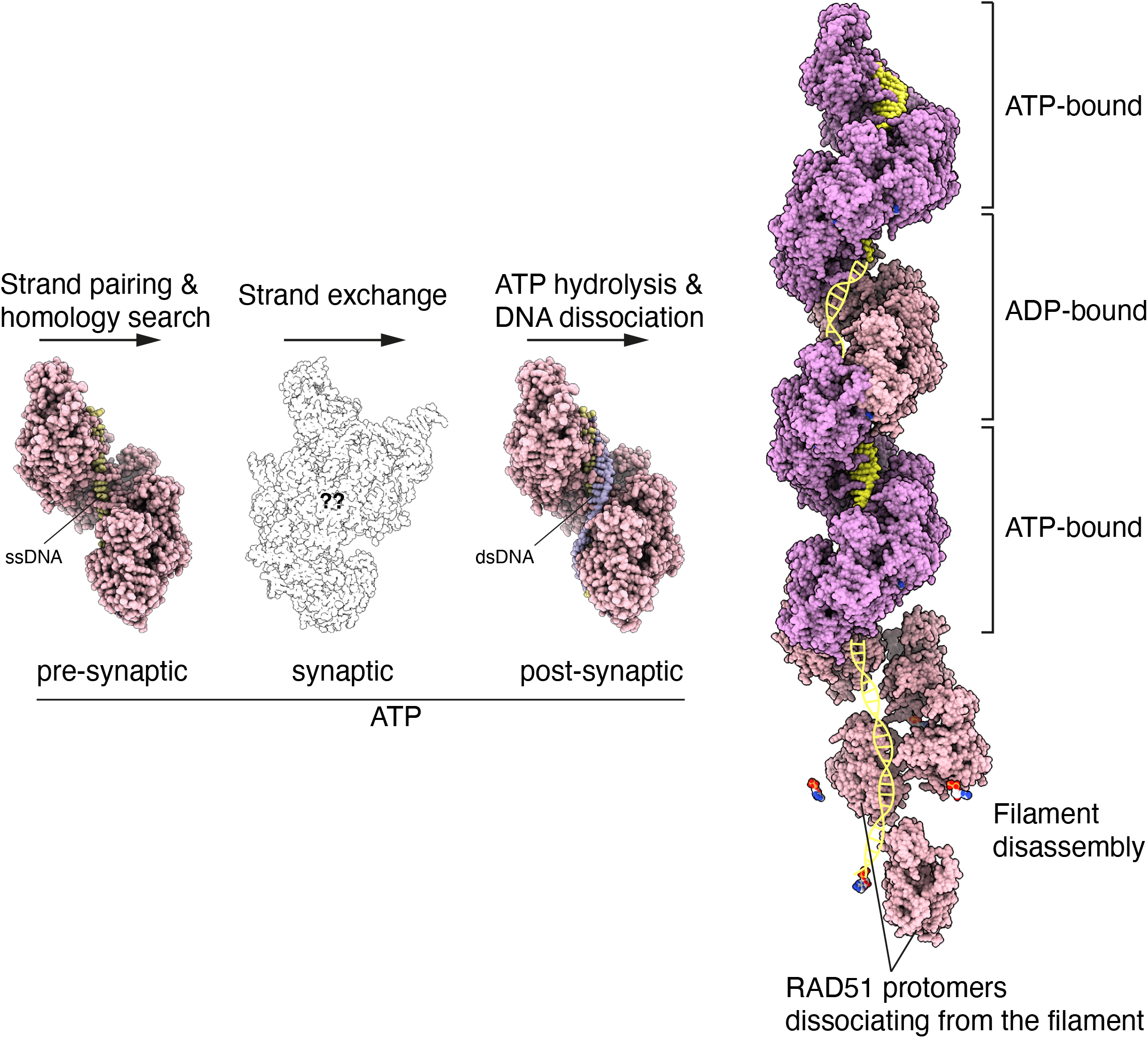
Structures of key RAD51 filament intermediates in the strand-exchange reaction. After completion of strand exchange, ATP hydrolysis at random points along the filament triggers dissociation of RAD51 from DNA, leading to segments of ATP-bound interspersed with DNA-free ADP-bound filament. ADP-bound RAD51 protomers that are licensed for disassembly dissociate from the filament end. RAD51 and DNA are shown as spacefill models, coloured pink for RAD51, pale yellow for ssDNA and pale yellow and mauve for the dsDNA strands. The pre-, post- and ADP-bound filaments correspond to cryoEM structures described in this work. The structure of the RAD51 synaptic filament is not known; an outline of the structure of bacterial RecA bound to a D-loop DNA is shown instead (PDB ID 7IY9).

The location and binding mode of an additional metal ion at the ATP site in human RAD51 NPFs resemble closely earlier reports of two potassium ions bound to the non-hydrolysable ATP analogue AMP-PNP in the crystallographic filament form of archaeal RadA(Wu et al., 2005). The interactions of one of the potassium ions with RadA and the nucleotide are very similar to the ones we observe for the second Ca^2+^ ion in the RAD51 filament. Indeed, the authors showed that two potassium ions observed in the crystallographic RadA filament can be replaced by one Ca^2+^ ion, with similar stimulatory effects on strand-exchange activity (Qian, He, Ma, et al., 2006). The promiscuity in the nature of the metal ion bound at the second site indicates that its function is to provide a positive charge that interacts favourably with the negatively charged C-end of L2’s *α*_289-295_ helix rather than a direct involvement in ATP hydrolysis, which would require a specific geometry of coordinating ligands. This argument is supported by the observation that a D316K mutation in human RAD51 stabilises the NPF and improves its recombinase function, and that the equivalent D302K mutation in archaeal RadA forms active NPFs in the absence of salt (Amunugama et al., 2012; Qian, He, Wu, et al., 2006).

Thus, the presence of the second metal ion in the RAD51 nucleoprotein filament confirms and extends to eukaryotic RAD51 the mechanism first proposed for how DNA binding by archaeal RadA is coupled to the hydrolysis state of the nucleotide: in the presence of ATP, the metal ion acts as a folding catalyst for the L2 loop, by helping specify its required conformation for DNA binding. This observation provides a structural rationale for the biochemical evidence that high concentrations of potassium and ammonium salts stimulate RAD51 activity (Liu et al., 2004; Stefan Sigurdsson et al., 2001). Conversely, loss of the gamma phosphate after ATP hydrolysis causes release of the second ion and impairment of DNA binding, as the L2 loop is no longer maintained in the correct conformation for interaction with DNA.

The cryoEM structure of the ADP-bound filament further shows that, rather than becoming disordered, the L2 loop takes up a new conformation that appears incompatible with DNA binding because of a steric clash with DNA. Thus, the identification of an ADP-bound filament intermediate that has dissociated from DNA provides a structural explanation for the requirement of ATP hydrolysis to licence the disassembly of the post-synaptic RAD51 filament (Figure 5). The increased conformational freedom acquired by the L2 loop upon release from its ATP-dependent DNA-binding conformation is consistent with a passive mode of DNA dissociation, in agreement with single-molecule observations that filament dissociation after ATP hydrolysis tends to be slow and incomplete (Hilario et al., 2009). As ATP hydrolysis takes places randomly throughout the filament (Tombline & Fishel, 2002), RAD51 dissociation from DNA is not cooperative; filament clearing from the DNA might be accelerated by accessory factors such as the human RAD54 translocase (Li et al., 2007; Mason et al., 2015; Solinger et al., 2002) or the worm helq-1 helicase (Ward et al., 2010). Alternatively, or in addition to licensing NPF disassembly, ATP hydrolysis might signal the completion of the strand-exchange reaction: in this model, random events of ATP hydrolysis along the filament would alter the L2 conformation and thus prevent further recombination, by making the filament architecture incompatible with the mechanism of strand-exchange.

Taking past and present evidence together, the metal ion-dependent mechanism for coupling ATP hydrolysis state to DNA binding and filament disassembly is now well-defined. However, an important question remains concerning the nature of the molecular determinants that prevent ATP hydrolysis within the filament until completion of strand-exchange and that might promote ATP hydrolysis afterwards. The cryoEM structures of pre- and post-synaptic filaments do not provide insight into this as they are essentially identical; indeed, dsDNA stimulates ATP hydrolysis by RAD51 to a lesser degree than ssDNA (Sung, 1994; Tombline & Fishel, 2002). As all aspects of RAD51 activity are subject to strict control in the cell, it is possible that the specific intervention of accessory factors might be required to repress ATP hydrolysis in the pre-synaptic filament and to stimulate it in the post-synaptic filament. The RAD51 paralogues have been reported to promote RAD51 filament activity in several ways, including filament assembly, remodelling and stabilisation of its active conformation (Belan et al., 2021; Roy et al., 2021; Taylor et al., 2015, 2016); however, no role in controlling the rate of ATP hydrolysis of the filament has yet been ascribed to them, although Xrcc2 has been shown to stimulate the ATPase activity of RAD51 (K. S. Shim et al., 2004). Further work will be required to clarify the mechanisms that regulate ATP hydrolysis within the RAD51 nucleoprotein filament.

## Acknowledgments

We would like to thank Joseph Maman for helpful comments on the manuscript, and Steve Hardwick for help with cryoEM data collection.

## Data availability

Coordinates and post-processed maps have been deposited in the PDB and EMDB databases, with accession codes 8BQ2 and 16170 (pre-synaptic filament), 8BR2 and 16197 (post-synaptic filament), 8BSC and 16224 (ADP-bound filament).

## MATERIALS & METHODS

### Expression and purification of human Rad51 protein

Calcium chloride-washed competent BL21 Rosetta *E*.*coli* cells were transformed with plasmid pMBP4-RAD51, containing the genes for human RAD51 and His_6_-MBP-BRCA2 BRC4. Transformants were selected by culturing on agar plates containing 50 mg/ml kanamycin. Transformants were resuspended in 2YT culture media and used to inoculate 1L cultures of 2YT culture media containing 50 mg/ml kanamycin. Cells were cultured at 37 °C until an OD_600_ of 0.7 was reached, before induction with 0.5 mM IPTG. Cultures were then incubated overnight at 15 °C before harvesting. Cell pellets were resuspended in resuspension buffer (500 mM NaCl, 50mM HEPES pH7.4) before flash freezing in liquid nitrogen for storage or used immediately for protein purification.

Cell pellets were defrosted and 1 tablet of sigma EDTA-free protease inhibitor cocktail was added per 50ml of resuspended cells. Cells were lysed by sonication on ice and lysate was centrifuged (40,000*g*, 1 hr, 4 °C) to remove cell debris. Lysate supernatant was loaded on a HisTrap HP 5 ml column preequilibrated with His Buffer A (20 mM HEPES pH 7.5, 300 mM NaCl, 5% glycerol) before isocratic elution in 40% His Buffer B (20 mM HEPES pH 7.5, 300 mM NaCl, 5 % glycerol, 500 mM imidazole). Eluted fractions were pooled and diluted in 1 ml batches by dropwise addition of a total of 3 ml of A125 Buffer (20 mM HEPES pH 7.5, 125 mM NaCl, 5 % glycerol, 2 mM DTT) while mixing. Diluted protein was loaded on a HiTrap Heparin 5 ml column preequilibrated with A125 buffer using a peristaltic pump at 4 °C, before a gradient elution of 12.5 % to 100 % buffer A1000 (20 mM HEPES pH 7.5, 1 M NaCl, 5 % glycerol, 2 mM DTT). Fractions containing RAD51 were concentrated to 2 ml using an Amicon 10,000 MWCO concentrator before purification by size exclusion using a Superdex200pg gel filtration column preequilibrated with RAD51 storage buffer (20 mM HEPES pH 7.5, 300 mM NaCl, 5 % Glycerol, 2 mM DTT). Eluted fractions containing RAD51 were pooled, concentrated to a final concentration of 85 *μ*M, snap frozen in liquid nitrogen and stored at -80 °C.

### Vitrification of RAD51 filament grids

#### Pre-synaptic filament

10 *μ*M RAD51 was incubated briefly on ice with a DNA 60mer (x60) in a buffer containing 20 mM TrisCl pH=7.5, 100 mM NaCl, 0.5 mM ATP, 5 mM CaCl_2_ prior to vitrification. The nucleotide sequence of the DNA was designed to avoid secondary structure propensity, using NUPACK (Zadeh et al., 2010). Cryo-EM grids were prepared by pipetting 3 µl of sample on a R1.2/1.3 Cu 300 mesh grid (Quantifoil) and plunge-freezing in liquid ethane using a Vitrobot Mark IV (ThermoFisher).

#### Post-synaptic and ADP-bound filaments

5 *μ*M RAD51 protein was incubated with DNA in a buffer containing 150 mM NaCl, 25 mM HEPES pH 7.5, 5 mM CaCl_2_, 2 mM DTT. The sample was supplemented with either 2 mM ATP or 2 mM ADP for the formation of post-synaptic- or ADP-containing filaments. For the preparation of post-synaptic filaments, the sample containing RAD51 was first incubated with 250 nM ssDNA-16SL for 15 minutes at 25 °C followed by incubation with 250 nM annealed R51-14 : R51-15 dsDNA for 15 minutes at 25 °C before vitrification. Glutaraldehyde was added to a final concentration of 0.05 % 3 minutes prior to grid freezing.

For the preparation of RAD51 ADP-bound filaments, annealed x60:cx60 dsDNA was added to the RAD51 reaction mix and incubated for 15 minutes at 25 °C prior to vitrification. UltrAuFoil R1.2/R1.3 300 mesh gold grids (Quantifoil) were glow-discharged twice (once on each side) for 1 minute using a PELCO easiGlow system (0.4 mBar, 30 mA, negative polarity). 6 *μ*l of reaction mix was applied to each grid before plunge-freezing in liquid ethane using a Vitrobot Mark IV robot (ThermoFisher), set to 100 % humidity, 4 °C, 2 second blot time and -8 blot force.

All DNA oligonucleotides were purchased PAGE-purified from IDT and resuspended to 100mM in TE buffer. DNA was annealed in TE buffer to a final concentration of 10 *μ*M dsDNA by boiling at 95 °C for 5 minutes followed by slow cooling to room temperature. All DNA sequences are listed here:

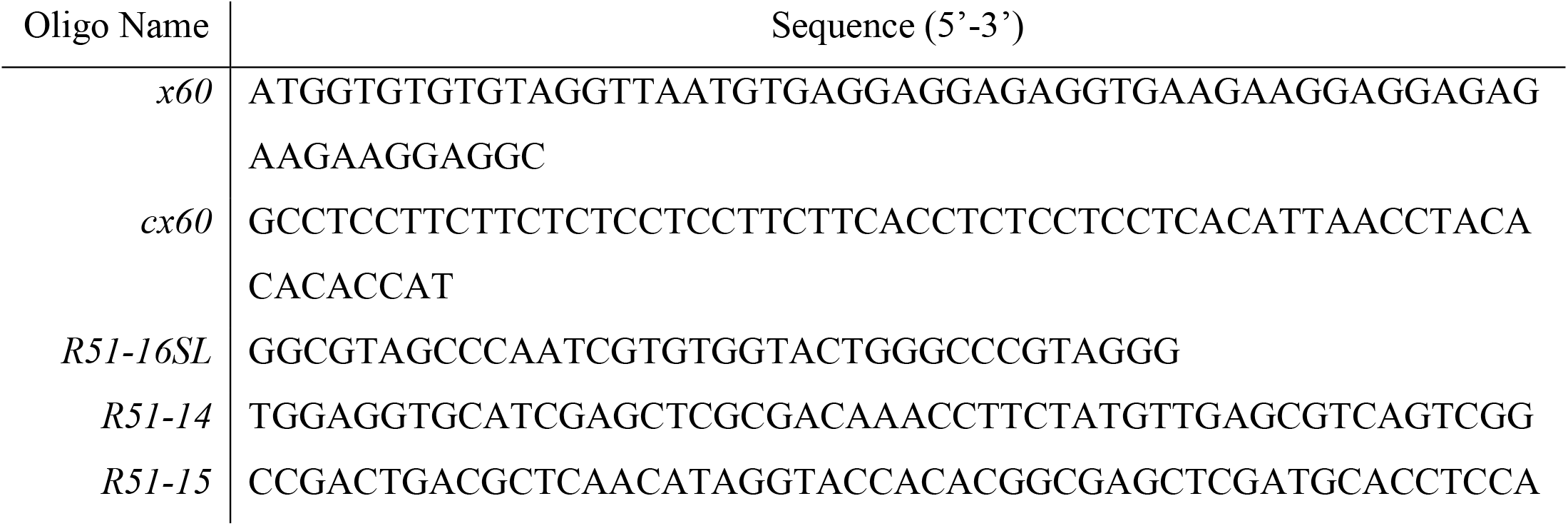

### CryoEM data collection and processing

Information about the data processing for the pre-, post- and ADP-bound filaments can be found in Supplementary table 1 and Supplementary figures 1-3.

#### Pre-synaptic filament

Data were collected on a Titan Krios microscope fitted with a Falcon III detector using the EPU package (FEI), at the Microscopy facility of the Nanoscience Centre. Data was processed using Relion 3.0 (Zivanov et al., 2018). Micrograph motion correction was performed using MotionCor2(Zheng et al., 2017) and CTF estimation was performed using CTFFIND4 (Rohou & Grigorieff, 2015). 850,217 particles were extracted using auto-picked coordinates and 723,142 particles were retained after rounds of 2D classification to remove badly aligned averages. An initial 3D model was generated from a subset of 5000 particles and subject to 3D classification. This initial 3D class was used for further 3D classification of the entire dataset, yielding three classes with very similar helical parameters. 3D class 2 with the best rotational accuracy was used as reference for 3D auto-refinement of the entire dataset, which converged at 4.2 Å. A second round of 3D auto-refinement after movie refinement and particle polishing improved the resolution to 4.1 Å. The map was further improved by masking and post-processing to a final resolution of 3.8 Å.

#### Post-synaptic and ADP-bound filaments

RAD51 grids were screened using a Talos Arctica microscope fitted with a Falcon III detector, and data were collected on a Titan Krios microscope fitted with a Gatan K3 detector at the Cryo-EM facility in the Department of Biochemistry. Data was processed using Relion 3.1 (Zivanov et al., 2018). Micrograph motion correction was performed using MotionCor2 (Zheng et al., 2017) and CTF estimation was performed using CTFFIND4 (Rohou & Grigorieff, 2015). Autopicking was performed based on manually picked 2D class averages with a picking threshold of 0.1. Autopicked particles were extracted with a box size of 260Å for the RAD51 postsynaptic filament and 200Å for the ADP-bound RAD51 filament and binned 4x and 2x respectively. 2D classification was performed iteratively on extracted particles with the option to ignore CTF correction until the first peak selected in addition to processing using ‘fast subsets’.

For the RAD51 postsynaptic filament, 543,102 high-resolution particles were identified following three rounds of single-particle 2D classification and were used to generate an initial 3D model. Single particle 3D classification produced highly anisotropic, poorly aligned classes, so particles were re-extracted and binned 4x and used to refine the initial 3D model, which was globally sharpened using Relion 3.1 to a final resolution of 2.9 Å. For the RAD51 ADP-bound filament, 62,657 high-resolution particles were identified following 3 rounds of 2D classification and were used to generate a 3D initial model. A first round of 3D classification using helical reconstruction (5 classes) generated three helical 3D models, all of which were unbound with DNA and exhibited helical parameters ranging from 52.9-56.0^°^ twist and 18.3-19.4 Å rise. 3D classification was repeated using one of the previous 3D classes as a starting model which generated four high resolution helical classes. These four 3D classes were extracted and binned 2x, yielding 40,481 particles which were used for refinement of the highest-resolution 3D class (class 2) to 4.9 Å, followed by global sharpening and density modification with *phenix*.*resolve_cryoem* to produce a map at 3.6 Å resolution.

### Model building and refinement

In all cases, RAD51-ATP protomers were individually fitted in the final filament maps, using the crystallographic coordinates of the human RAD51-ATP structure in filament form (PDB ID 5NWL) (Brouwer et al., 2018). For the pre-synaptic NPF, a ssDNA 30mer consisting of a tandem repeat of three GGA nucleotides was generated in Coot (Emsley & Cowtan, 2004) and fitted in the map. GGA was chosen as the consensus RAD51-binding site, based on the nucleotide frequency in the 20 RAD51-binding sites present in the ssDNA 60mer. For the post-synaptic NPF, a double-stranded DNA sequence corresponding to a fully-complementary portion of the R51-14 : R51-15 duplex was fitted in the map.

Fitting of the filament models to the map was improved using real-space refinement as implemented in Phenix (Adams et al., 2010). For all filament models, the two protomers occupying the outermost positions in the map were left out of the final refined model as they had poorer model-to-map correlation coefficients.

### Electrophoretic mobility shift assay (EMSA)

EMSA reactions were prepared in buffer: 25 mM HEPES pH 7.5, 150 mM NaCl, 2 mM DTT, 5 mM CaCl_2_, supplemented with either 2 mM ATP or a titration of 0.5, 1, 2, 5, 10 mM ADP. Fluorescein-labelled dx60:cdx60 dsDNA was added to a final concentration of 500 nM and incubated with RAD51 at a final concentration of 10 *μ*M at 25 °C for 15 minutes prior to loading. Immediately before loading, samples were mixed with a 0.5 : 1 volume of 50 % (w/v) sucrose solution. Samples were run on a pre-run 0.5 % agarose gel in 0.5x TB buffer (45 mM Tris, 45 mM boric acid) for 2.5hr at 4^°^C and 35V. The gel was imaged using a Typhoon FLA9000 by excitation at 488 nm.

