## Supplementary information for "A metal ion-dependent mechanism of RAD51 nucleoprotein filament disassembly"

**SUPPLEMENTARY FIGURE LEGENDS**

**Supplementary Figure 1.** CryoEM data processing for the pre-synaptic RAD51 NPF. **A** Representative micrograph of RAD51 pre-synaptic filament particles. **B** Representative 2D average class. **C** 3D classification and refinement. **D** Fourier Shell Correlation between half maps (top) and model to map (bottom). The resolution at 0.143 FSC is indicated by the intersection of the dashed lines.

**Supplementary Figure 2.** CryoEM data processing for the post-synaptic RAD51 NPF. **A** Representative micrograph of RAD51 post-synaptic filament particles. **B** Representative 2D average classes. **C** 3D classification and refinement. **D** Fourier Shell Correlation between half maps (top) and model to map (bottom). The resolution at 0.143 FSC is indicated by the intersection of the dashed lines.

**Supplementary Figure 3.** Biochemical preparation and cryoEM data processing of the ADP-bound RAD51 filament. **A** Electrophoretic mobility shift assay of dsDNA in the presence of RAD51 and increasing amounts of ADP (see Methods for details). **B** Representative micrograph of RAD51 post-synaptic filament particles. **C** Representative 2D average classes. **D** Initial 3D model, two rounds of 3D classification and 3D refinement. **E** Fourier Shell Correlation between half maps (top) and model to map (bottom). The resolution at 0.143 FSC is indicated by the intersection of the dashed lines.

**Supplementary Figure 4.** L2 loop conformation in the ADP-bound RAD51 filament. The L2 segment not modelled as not visible in the cryoEM reconstruction is drawn as yellow dots. **A** Two adjacent RAD51 protomers are shown in light brown and orange. The L2 loop is highlighted in yellow, with residue side chains shown. Residues that are engaged in prominent hydrophobic interactions are labelled. **B** View of the L2 loop conformation that highlights its likely steric clash with the DNA. Colour-coding as in panel A. The inner strand of the dsDNA from the post-synaptic filament is shown as stick model and transparent molecular surface.

**Supplementary Figure 5.** Evidence for the presence of a second metal ion at the ATP-binding site in high-resolution cryoEM maps of pre-synaptic human RAD51-AMPPNP-$Mg^{2+}$ (EMD 31158, PDB 7EJC), pre-synaptic human DMC1-AMPPNP-$Ca^{2+}$ (EMD 30311, PDB 7C9C) and post-synaptic yeast DMC1-ATP-$Mg^{2+}$ (EMD 31154, PDB 7EJ7). The position of the density peak attributable to the second metal cation is indicated by an arrow. The ATP and the side chains of important ATP binding-residues are drawn explicitly.

**Supplementary Table 1. CryoEM data collection and real-space refinement.**

*Data Collection*

| | **Pre-synaptic** | | **Post-synaptic** | | **ADP-bound** | |
|---|---|---|---|---|---|---|
| Microscope | Titan Krios | | | | | |
| Voltage (keV) | 300 | | | | | |
| Detector | Falcon III | | K3 (Gatan) | | | |
| Collection mode | Integration | | Counting | | | |
| Magnification | 59,000x | | 130,000x | | | |
| Defocus range (μm) | -3.0 to -1.5 (0.3) | | -2.5 to -0.9 (0.2) | | | |
| Num. movies | 1339 | | 3851 | | 9214 | |
| Frames/movie | 38 | | 40 | | 50 | |
| Pixel size (Å/pixel) | 1.43 | | 0.652 (super-resolution 0.326) | | | |
| Electron dose (e$^-$/Å$^2$/s) | 33 | | 47.22 | | 56.36 | |
| Exposure (s) | 2 | | 1.61 | | 1.34 | |
| Picked particles | 850,217 | | 1,051,037 | | 1,527,085 | |
| Final particles | 723,142 | | 543,102 | | 40,481 | |
| Processing method | Helical reconstruction | | Single particle | | Helical reconstruction | |
| Map resolution: | masked | unmasked | masked | unmasked | masked | unmasked |
| $d_{FSC}$ half maps, 0.143 | 3.7 | 3.5 | 2.8 | 3.0 | 4.2 | 4.5 |
| $d_{FSC}$ model, 0.143 | 3.7 | 3.7 | 2.6 | 2.7 | 3.7 | 3.9 |
| Helical twist (°) | 56.7 | | - | | 53.1 | |
| Helical rise (Å) | 16.6 | | - | | 18.6 | |

*Real-space refinement*

| | Pre-synaptic | Post-synaptic | ADP-bound |
|---|---|---|---|
| Composition: | | | |
| Non-H atoms | 22331 | 15574 | 16982 |
| Residues: | | | |
| Protein | 2790 | 1866 | 2191 |
| Nucleotide | 30 | 40 | - |
| Water | - | 120 | - |
| Ligands: | | | |
| ATP | 9 | 6 | |
| ADP | | | 7 |
| $Ca^{2+}$ | 18 | 12 | 7 |
| Correlation coefficients:[1] | | | |
| CC, mask | 0.86 | 0.83 | 0.69 |
| CC, peaks | 0.66 | 0.66 | 0.52 |
| CC, volume | 0.85 | 0.78 | 0.68 |
| <CC>, ligands | 0.86 | 0.70 | 0.65 |
| Bonds (rmsd): | | | |
| Length(Å) | 0.003 | 0.012 | 0.005 |
| Angles (°) | 0.542 | 0.643 | 0.601 |
| MolProbity score[2] | 1.42 | 1.94 | 2.25 |
| Clash score | 7.62 | 16.40 | 15.99 |
| Ramachandran plot (%): | | | |
| Outliers | 0.00 | 0.00 | 1.29 |
| Allowed | 1.96 | 2.93 | 6.47 |
| Favoured | 98.04 | 97.07 | 92.23 |
| Rotamer outliers (%) | 0.00 | 1.20 | 1.20 |
| Cb outliers (%) | 0.00 | 0.00 | 0.00 |
| ADP (B-factors), mean: | | | |
| Protein | 67.36 | 27.54 | 36.07 |
| Nucleotide | 58.40 | 79.69 | - |
| Ligand | 53.90 | 20.34 | 42.73 |
| Water | - | 31.05 | - |

1. Afonine, P.V., Klaholz, B.P., Moriarty, N.W., Poon, B.K., Sobolev, O.V., Terwilliger, T.C., Adams, P.D., and Urzhumtsev, A. (2018). New tools for the analysis and validation of cryo-EM maps and atomic models. Acta Crystallographica Section D: Structural Biology *74*, 814–840. 10.1107/s2059798318009324.

2. Chen, V.B., Arendall, W.B., Headd, J.J., Keedy, D.A., Immormino, R.M., Kapral, G.J., Murray, L.W., Richardson, J.S., and Richardson, D.C. (2010). MolProbity: all-atom structure validation for macromolecular crystallography. Acta Crystallogr Sect D Biological Crystallogr *66*, 12–21. 10.1107/s0907444909042073.

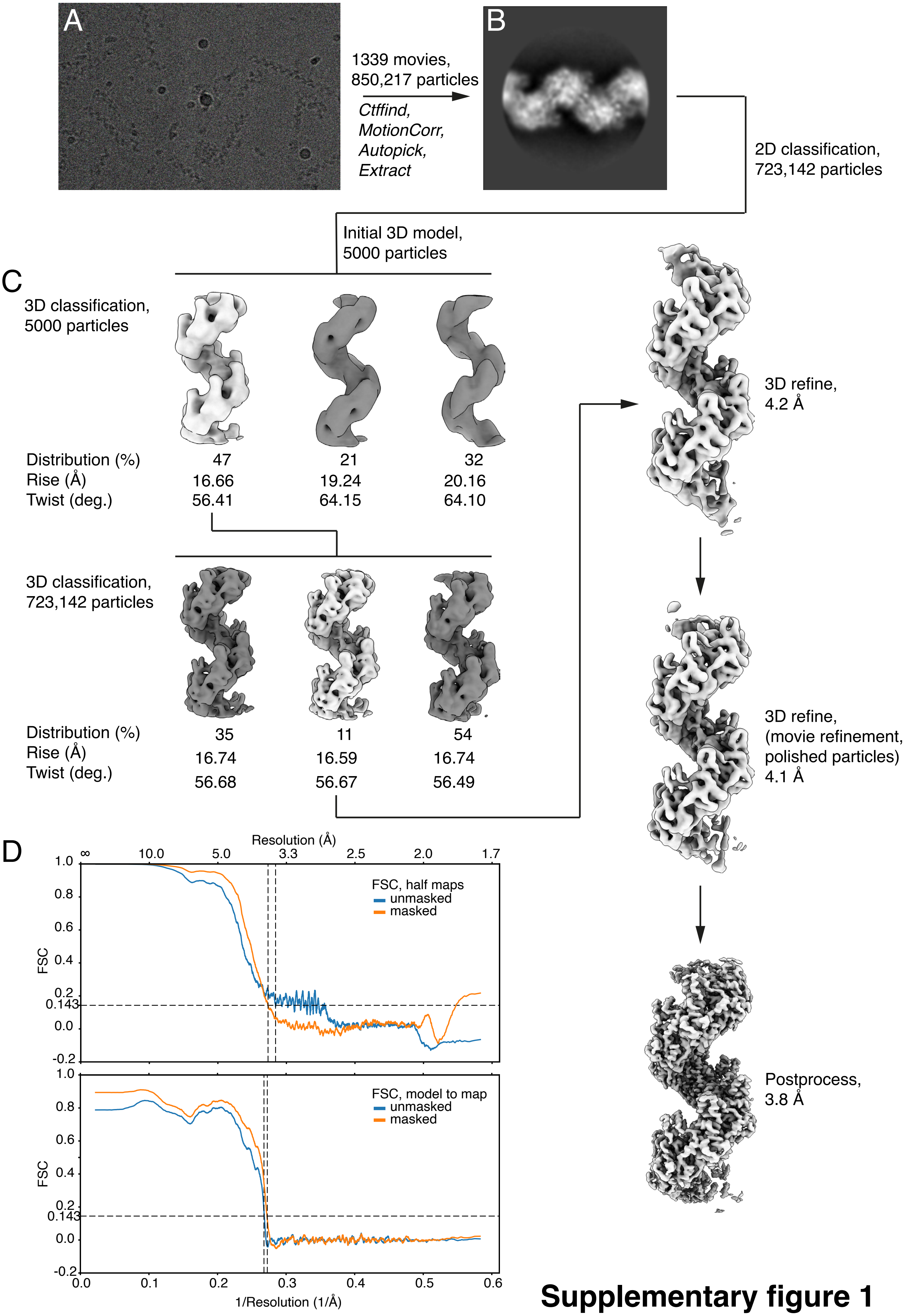


**Supplementary figure 1**

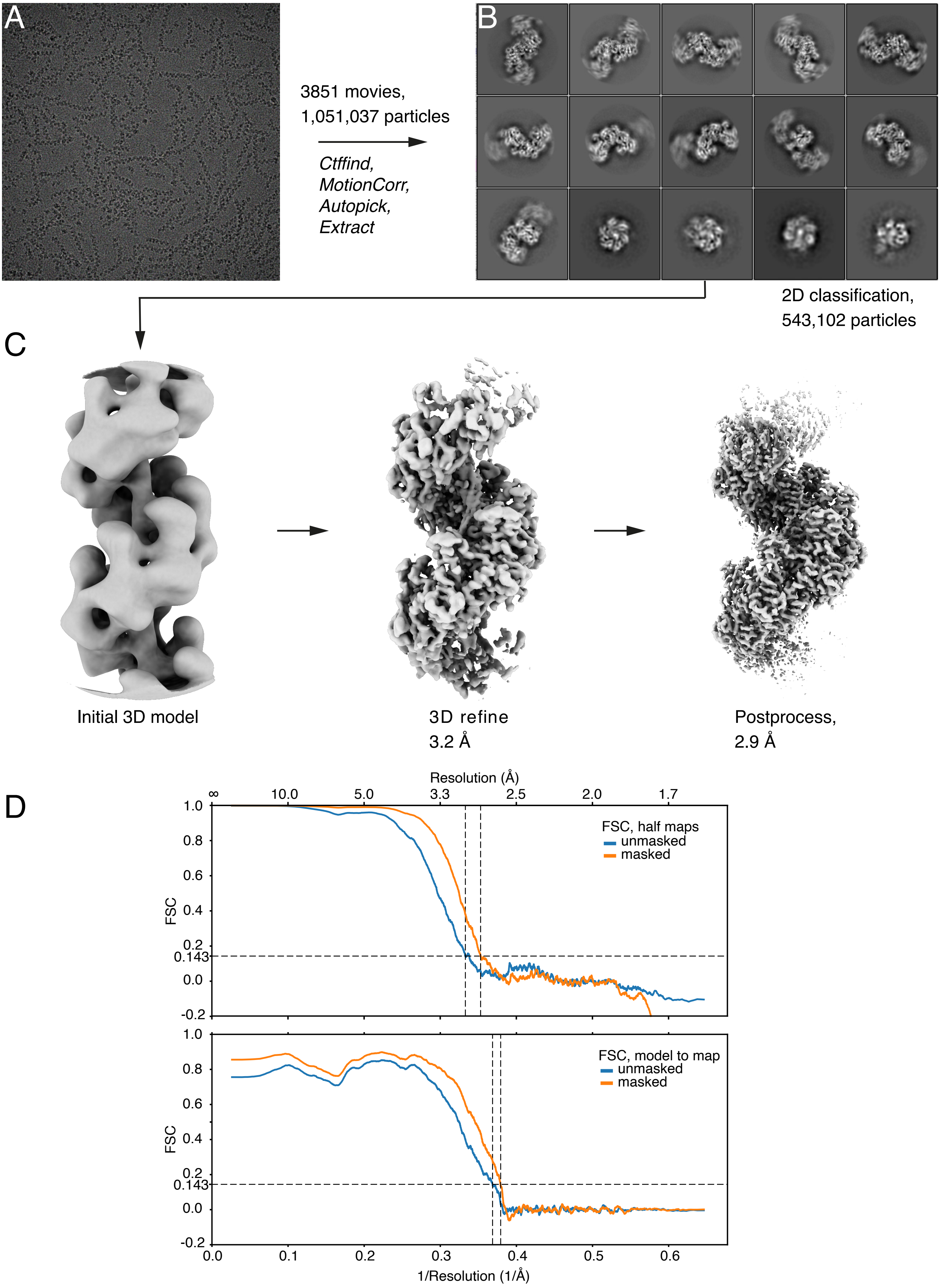


**Supplementary figure 2**

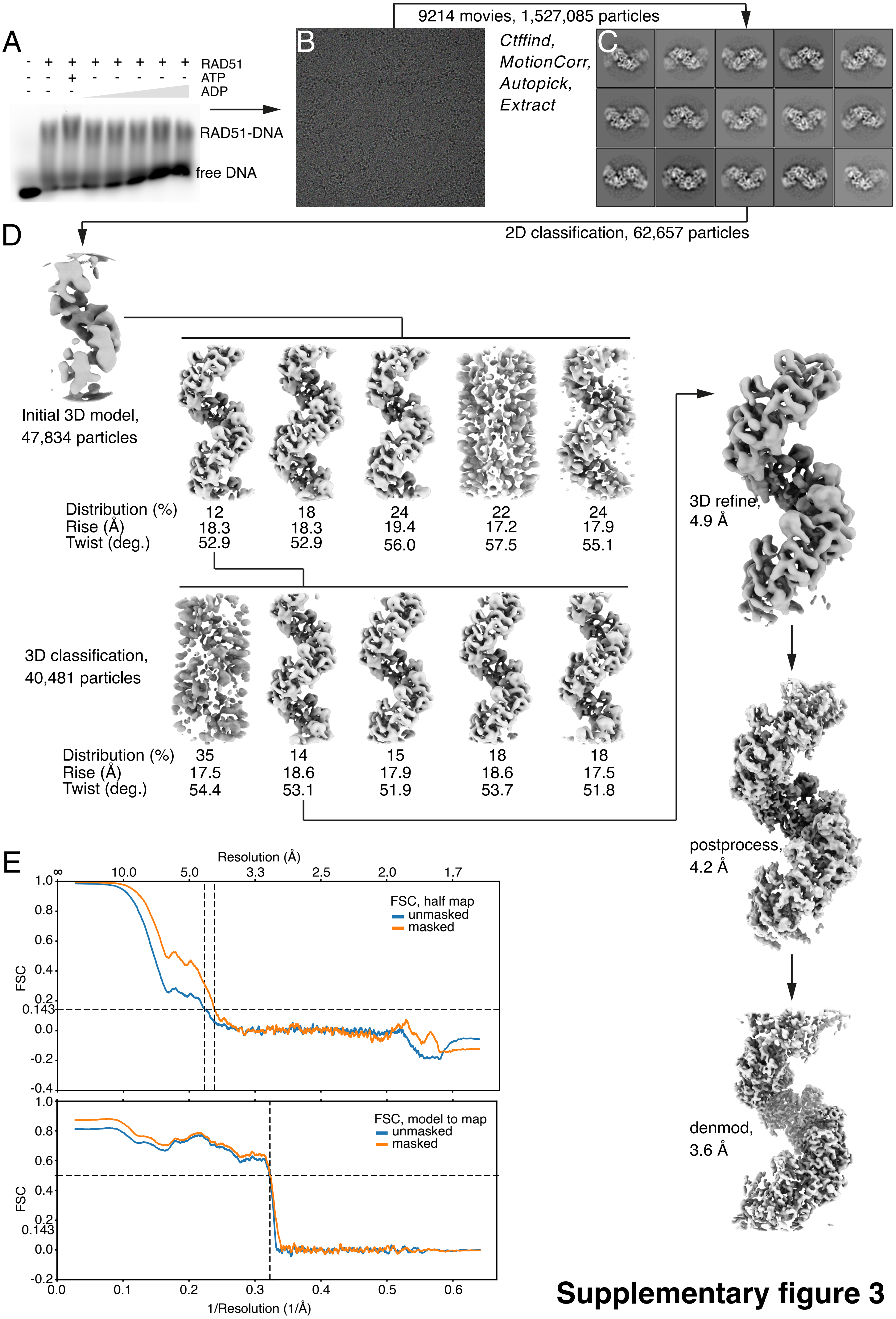


**Supplementary figure 3**

A

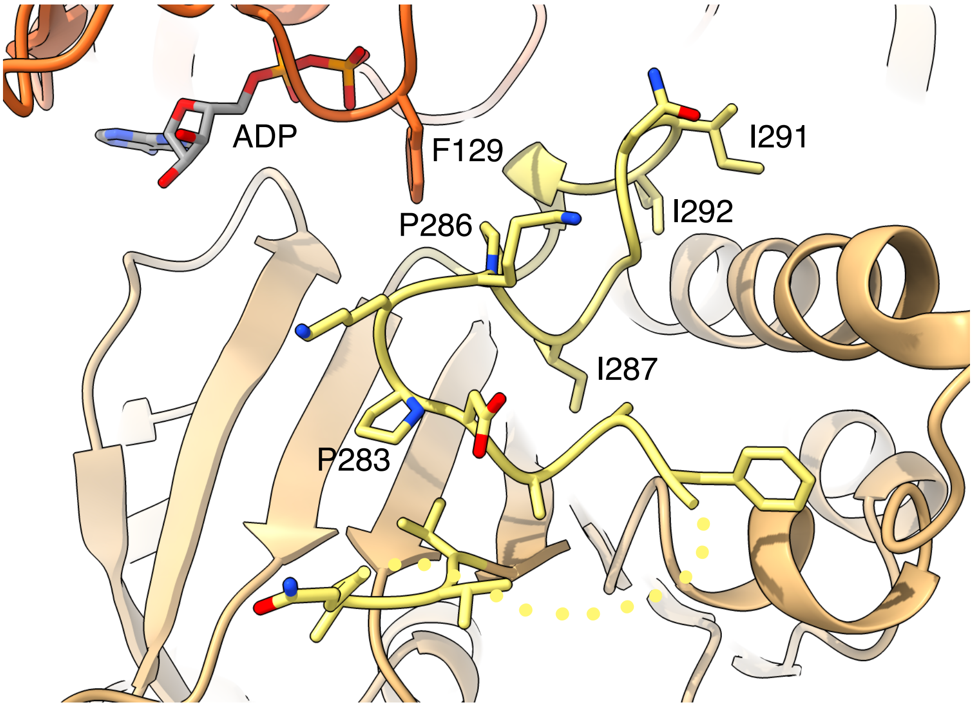


B

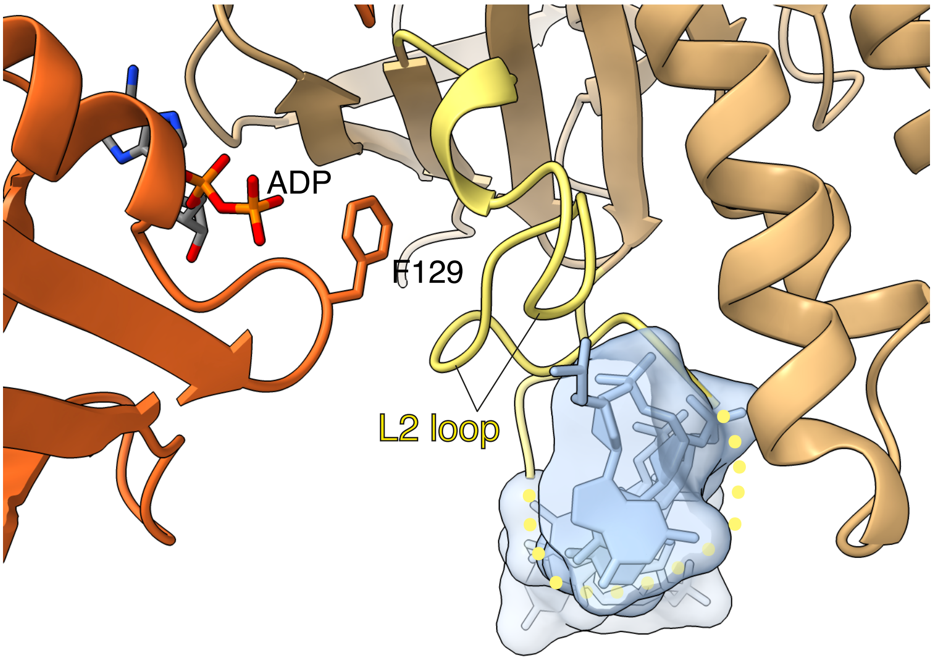


**Supplementary figure 4**

RAD51
Pre-synaptic filament
PDB ID: 7EJC
2.97 Å

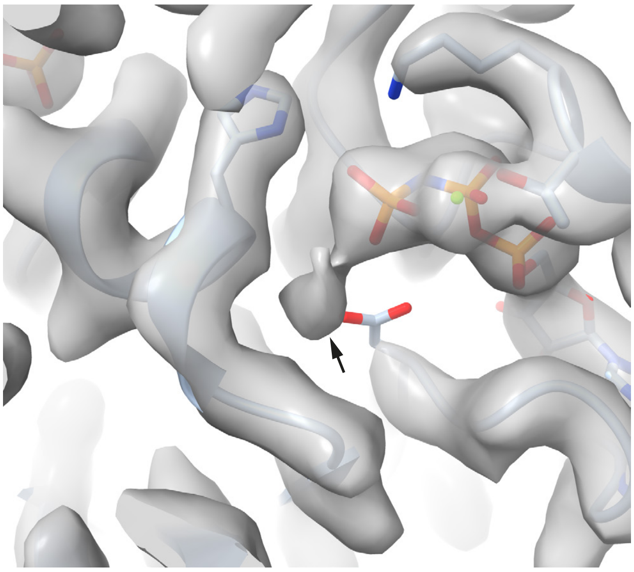

DMC1
Pre-synaptic filament
PDB ID: 7C9C
3.33 Å

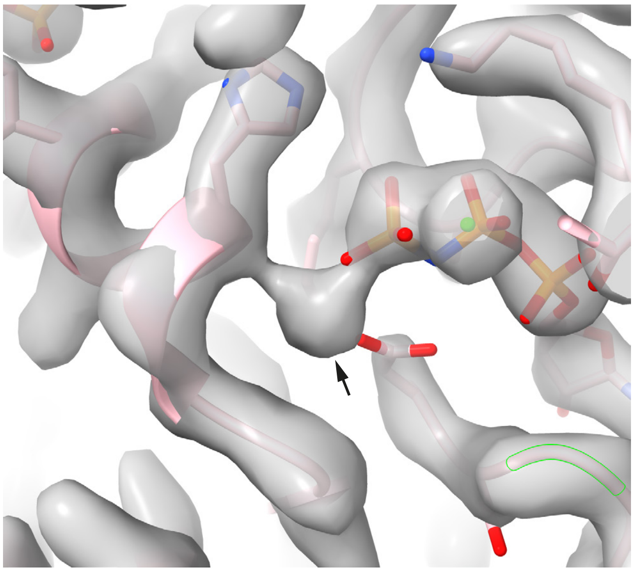

DMC1
Post-synaptic filament
PDB ID: 7EJ7
3.41 Å

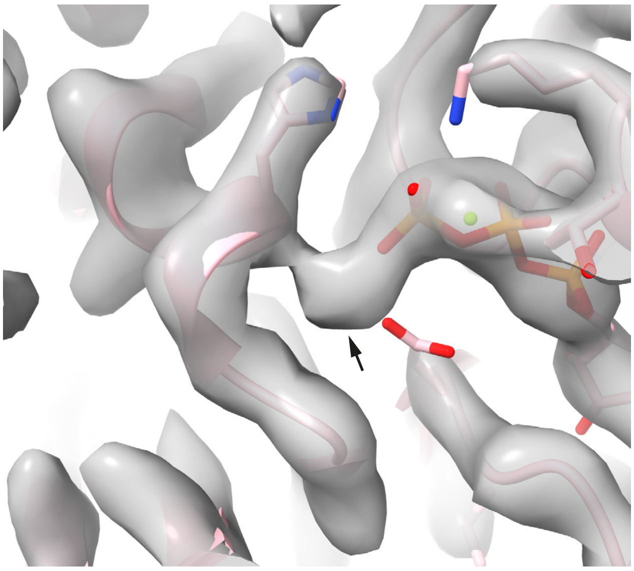

**Supplementary figure 5**
